# Ziqi Dihuang decoction ameliorates thrombosis in septic rats by inhitbiting plasminogen activator inhibitor-1

**DOI:** 10.1101/2023.04.05.535767

**Authors:** YanXia Geng, ShuYe Fei, YingHao Pei, QiuHua Chen, Jian Wang, Hua Jiang

## Abstract

**Background and Aim:** Sepsis is now a global medical burden with high morbility and mortality. The focus of this study was to observe and elaborate the effect of Ziqi Dihuang (ZQDH) decoction on inflammatory and thrombosis-related parameters in septic rats.

**Experimental procedure:** A rat model of sepsis was established by cecal ligation and puncture (CLP). 24 male Sprague-Dawley (SD) rats were randomly divided into Sham group, CLP group, ZQDH-1ow group (0.735g/kg) and ZQDH-high group (1.47g/kg). Rats in ZQDH groups were given ZQDH decoction by gavage for 7 days before CLP, while rats in Sham and CLP groups were given the same amount of normal saline. Leukocytes and percentage of neutrophils (N %) in blood, and inflammatory cell infiltration of liver, kidney and lung were used to assess systemic inflammatory response. Coagulation and fibrinolytic indexes included platelet count, coagulation function, fibrin deposition, and levels of tissue plasminogen activator (tPA) and plasminogen activator inhibitor-1 (PAI-1) in serum, liver, kidney and lung.

**Results:** ZQDH decoctioninhibited N% growth in blood and inflammatory cell infiltration in the lung of CLP rats. Moreover, ZQDH decoctionalso ameliorated thrombocytopenia and prothrombin time, alleviated renal fibrin deposition, and improved tPA and PAI-1 levels in kidney.

**Conclusion:** These results suggest that ZQDH decoction can dose-dependently relieve systemic inflammatory injury and regulate coagulation-fibrinolysis system in septic rats, which may be mediated by PAI-1.

**Highlights of the findings and novelties:**

ZQDH decoction relieves systemic inflammatory injury in sepsis;
ZQDH decoction regulates coagulation-fibrinolysis system in sepsis;
ZQDH decoction inhibits fibrinolysis by reducing inhibitor-1 (PAI-1) level in sepsis.

## 1. Introduction

Sepsis is an acute syndrome triggered by infection that can lead to life-threatening organ dysfunction ^[1]^. With an annual incidence above 30 million cases and an acute mortality rate about 26.0%, sepsis has become a global burden, and researchers have been struggling to develop new treatments for it ^[2]^. Sepsis is often accompanied by coagulation and fibrinolytic dysfunction, which puts patients at risk for both blood clots and bleeding, manifested as thrombocytopenia, decreased fibrinogen, increased fibrinogen degradation products and microvascular thrombosis, which can lead to disseminated intravascular coagulation (DIC). Plasminogen activator inhibitor −1 (PAI-1) is an inhibitor of fibrinolysis, and an important element of the imbalance between clot formation and fibrinolysis ^[3]^. As a risk factor for thrombosis, it is also known to be an considerable predictor of severity and mortality in sepsis ^[4–5]^.

Sepsis-induced DIC is characterized by inhibition of fibrinolysis and is prone to develop into multiple organ dysfunctions. At present, there is no definite treatment for it ^[6]^. In traditional Chinese medicine, sepsis belongs to the category of febrile diseases. The theory of blood stasis and heat stasis of Chinese medical master Zhou Zhongying provides a new idea for the treatment of sepsis-related coagulopathy by cooling blood and dispersing blood stasis ^[7]^. In the Ziqi Dihuang (ZQDH) decoction of this study,

Rhubarb, Lithospermum and Radix Paeoniae Rubra are typical traditional Chinese medicine for clearing heat and cooling blood, promoting blood circulation and removing blood stasis. We intend to investigate the effect of ZQDH decoction on sepsis-induced coagulopathy and thrombosis in a rat model of cecal ligation and puncture (CLP).

## 2. Materials and methods

### 2.1 Drugs and chemicals

ZQDH decoction consists of five herb granules, including Rehmanniae (Di Huang, 30g), Paeoniae Rubra (Chi Shao, 10g), Lithospermum (Zi Cao, 10g), Panax notoginseng (San Qi, 10g) and Rhubarb (Da Huang, 10g). All the herb granules were purchased from Tianjiang Pharmaceutical Co Ltd, Taizhou, China. Enzyme-linked immunosorbent assays (ELISA) kits for tPA and PAI-1 were obtained from Shanghai Enzyme-linked Biotechnology Co., Ltd. (Shanghai, China).

### 2.2 Animals and experimental protocol

A total of 24 male Sprague-Dawley rats, weighing 200 ±20 g and 8-10 weeks old, were provided by the Animal Laboratory of Jiangsu Province Hospital of Chinese Medicine, with animal study approval number of SCXK (Jing) 2019-0010. Rats were housed in cages with a temperature of 20 ±1°C and relative humidity of 45%, and had free access to tap water and standard laboratory feed. Rats were randomly divided into four groups: Sham group (n=6), CLP group (n=6), ZQDH-low (7.35g/kg, n=6) and ZQDH-high group (14.7g/kg, n=6).The normal dosefor rats (7.35g/kg.d) was determined based on the normal dose for adults (70g/d), and thebody surface area ratio between adults and rats. In this study, the normal dose was referred to as “low dose” and twice the normal dose was referred to as “high dose”. ZQDH herb granules were dissolved in distilled water to a final concentration of 0.735g/ml or 1.47g/ml, and were administered to rats by oral gavage for 7 days before CLP operation. Rats in Sham and CLP groups were gavaged with the same volume of normal saline. All procedures were implemented in accordance with the Animal Management Regulations of the Ministry of Health of China.

Septic rat model was established by cecal ligation and puncture (CLP) surgery^[8]^. Rats were fasted and given free access to water for 12 h prior to CLP surgery. After anesthetized with 1% pentobarbital sodium (40mg/kg, ip), rats were subjected to a midline incision of ~2 cm on the anterior abdomen. The cecum was carefully separated, exteriorized, and ligated by 3-0 suture at half of the cecum. Then ligated cecum was punctured using a 12G needle to create two holes and was gently squeezed to discharge a small amount of fecal content. Thereafter, put the cecum back into the abdominal cavity, suture and close the abdomen. Rats in the Sham group were subjected to identical abdominal incision and intestinal manipulation, but the cecum was neither ligated nor punctured. All surgical procedures were completed within 15 minutes. All rats were subcutaneously injected with normal saline (0.04ml/g) for fluid resuscitation and were allowed free access to food and water after surgery.

All rats were sacrificed 12 hours after operation. Blood samples were collected by heart puncture, and tissue specimens (liver, kidney and lung) were preserved for subsequent experiments.

### 2.3 Histopathological testing

Tissue specimens (liver, kidney and lung) were fixed with 4% paraformaldehyde at 4°C for >24 h, embedded in paraffin and serially sectioned (5 μm). Hematoxylin-eosin (HE) and Martius-Scarlet-Blue (MSB) were used to quantify the inflammatory cells infiltration and fibrin deposition, respectively. For each assessment, 40× high power field images of affected hepatic lobules, glomeruli, or alveoli were captured and 10 images from each group were randomly selected. Image J software was used for image process and analysis. The number of red pixels corresponding to fibrin in MSB image was automatically calculated using the threshold color plug-in and represented as a histogram.

### 2.4 Blood test of inflammatory and coagulation indexes

White blood cell count (WBC), percentage of neutrophils (N %), platelet count (PLT) and mean platelet volume (MPV) were tested using an automatic haematology analyzer (SysmexXS-800i, Japan). Prothrombin time (PT), activated partial thromboplastin time (APTT), and fibrinogen (FIB) were detected with an automatic coagulation analyzer (STAGOSTA-R MAX, France).

### 2.5 Measurement of tPA and PAI-1

The contents of tPA and PAI-1 in serum and tissues (liver, kidney and lung) in each group were detected with ELISA kit. The serum samples were obtained by centrifugation of blood samples at 3,000 g for 15 min at 4°C. The tissue supernatant was obtained by centrifuging the tissue homogenate at 5,000 g for 15 min at 4°C. All procedures were carried out according to the manufacturer’s protocol.

### 2.6 Statistical analysis

SPSS software (version 26.0, IBM Corp) was used for data analysis. All continuous data were presented as the mean ± standard deviation and analyzed using one-way ANOVA. Pairwise comparisons were performed with LSD post-hoc test when equal variances assumed, or Tamhane T2 post-hoc test when equal variances not assumed. A *p* value less than 0.05 was considered statistically significant.

## 3. Results

### 3.1 HE staining of liver, kidney and lung

As shown in Figure 1, there was no significant difference in the infiltration of inflammatory cells in liver and kidney among the four groups of rats. CLP only caused swelling and necrosis of some hepatocytes and infiltration of a few inflammatory cells. In the kidney, CLP induced increase of glomerular interstitial, capillary congestion, interstitial edema and some inflammatory cell infiltration. In the lung, however, the alveolar wall was notably thickened and a large number of inflammatory cells infiltrated into the interstitium after CLP. The infiltration of inflammatory cell was significantly alleviated in ZQDH treatment groups, in a dose-dependent manner.

**Figure 1.**
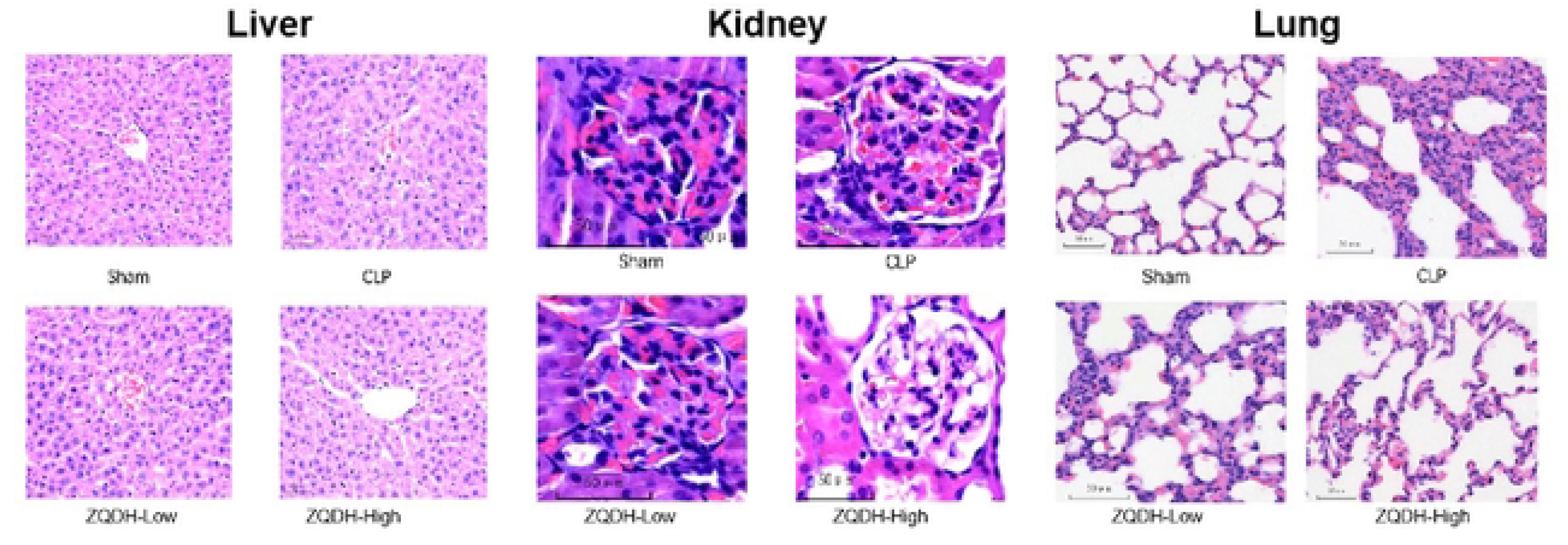
HE staining of liver, kidney and lung. Pulmanary inflammatory cells infiltration were significantly reduced in ZQDH groups than the CLP group, in a dose-dependent manner.

### 3.2 MSB staining of liver, kidney and lung

As shown in Figure 2, there was no significant difference in liver MSB staining among four groups of rats. The fibrin deposition volume in kidney and lung of CLP group were considerably larger than that of Sham group. With ZQDH treatment, the fibrin deposition in glomeruli was significantly reduced, and it was further attenuated in ZQDH-high group than ZQDH-low group. But this effect was not seen in the lung.

**Figure 2.**
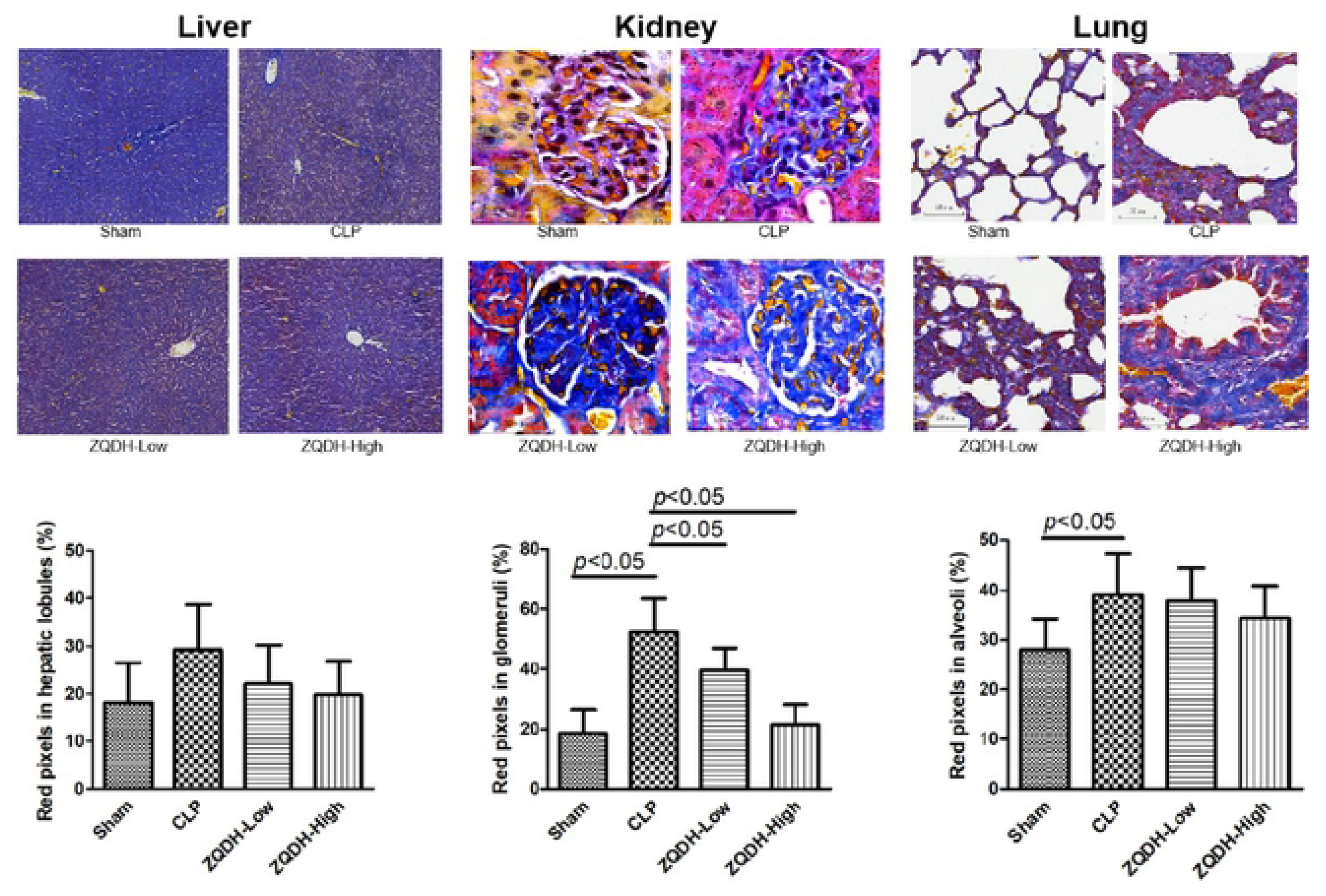
MSB staining and relative volume fraction of fibrin of liver, kidney and lung. Fibrin deposits are red-colored, and the area of renal fibrin deposition was significantly lower in ZQDH groups than CLP group in a dose-dependent manner.

### 3.3 Leukocyte and platelet in the blood

No significant difference was seen in WBC and MPV among four groups of rats (Fig. 3A& 3D). As shown in Figure 3B, the N% level in blood increased significantly after CLP and decreased with ZQDH intervention. Similarity, the number of PLT in CLP group was noticeably lower than the Sham group, and it was improved after high dose of ZQDH treatment (Fig. 3C).

**Figure 3.**
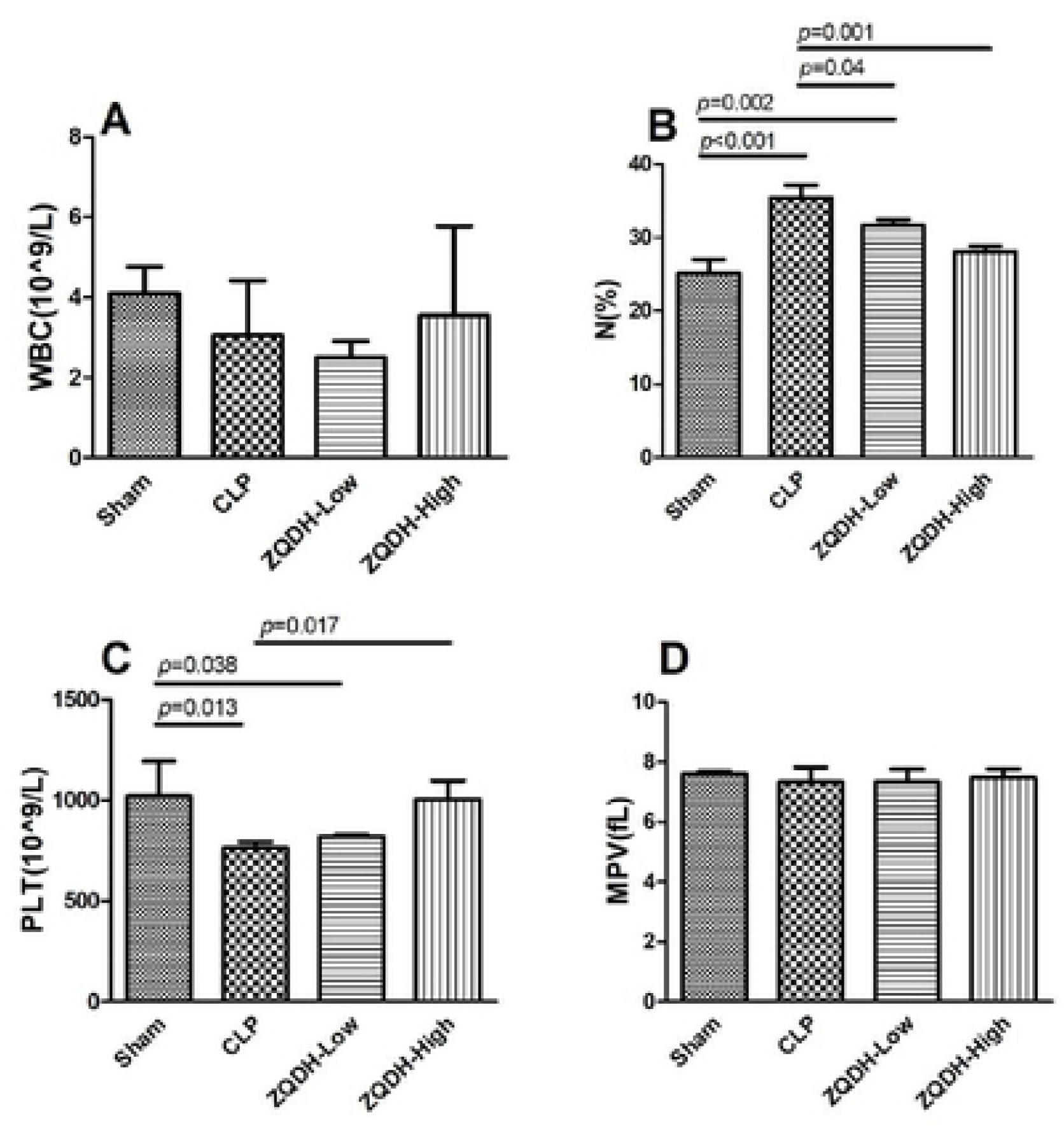
Concentrations of white blood cell count (WBC, Figure 3A), percentage of neutrophils (N%, Figure 3B), platelet count (PLT, Figure 3C) and mean platelet volume (MPV, Figure 3D) in the blood. Levels of N% and PLT were significantly improved in ZQDH group when compared with the CLP group.

### 3.4 Serum coagulation function indexes

As shown in Figure 4, no significant difference was observed in APTT among the four groups. Levels of PT and FIB were markedly decreased in CLP group than those in Sham group. After low or high dosage of ZQDH intervention, serum PT level elevated, while FIB did not.

**Figure 4.**
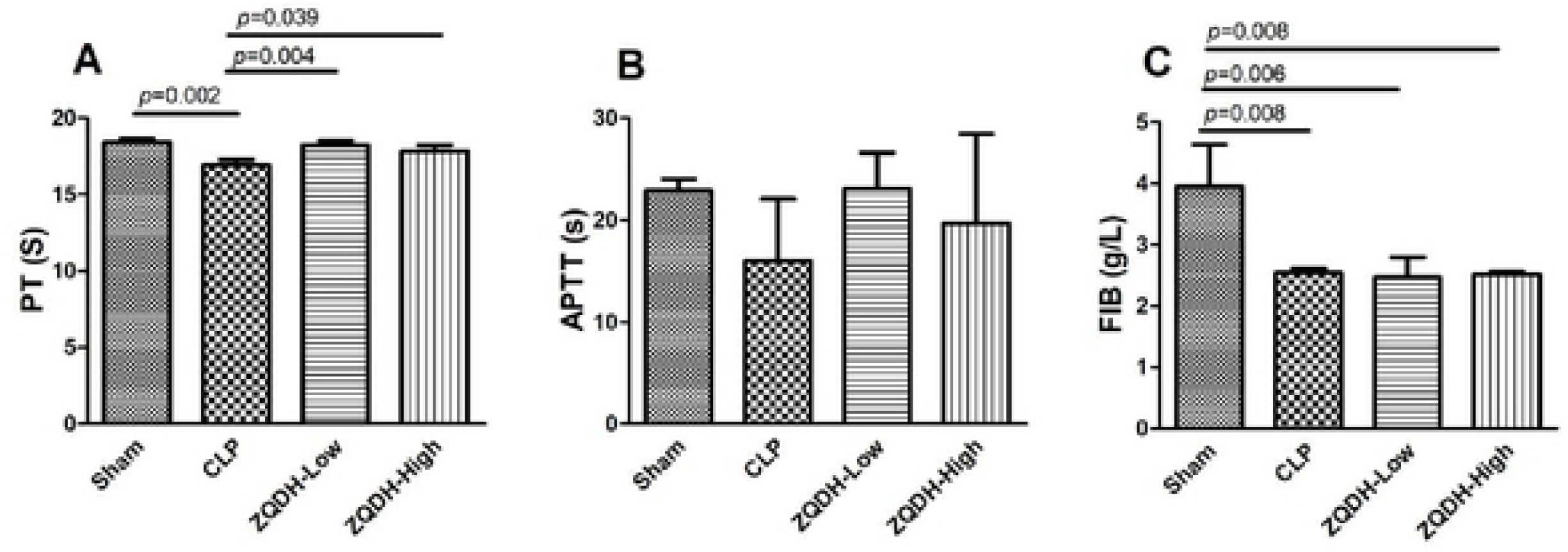
Serum levels of prothrombin time (PT), activated partial thromboplastin time (ΛPTT), and fibrinogen (FIB). PT and FIB levels in CLP group were significantly decreased, and the level of PT was improved after ZQDH intervention, but not in FIB.

### 3.5 tPA and PAI-1 in serum, liver, kidney and lung

Figure 5 shows that serum level of tPA and PAI-1 did not differ significantly among four groups of rats. CLP obviously increased level of PAI-1 in liver and kidney, as well as tPA in liver, kidney, and lung. With ZQDH intervention, concentrations of tPA in kidney and lung, and PAI-1 in liver and kidney were significantly decreased in the ZQDH-high group.

**Figure 5.**
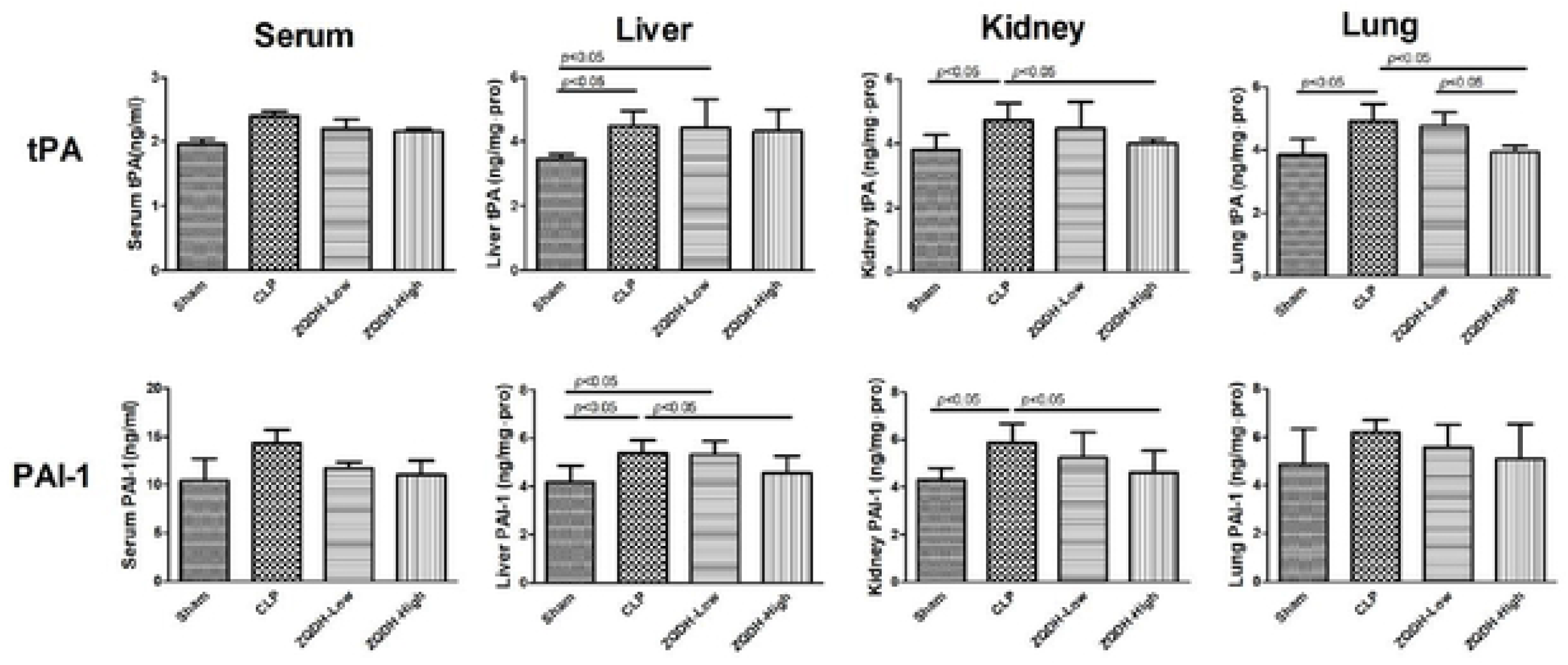
Concentrations of tissue plasminogen activator (tPA) and plasminogen activator inhibitor-1 (PAI-1) in serum and tissues of rats in four groups.

## 4. Discussion

Cecal ligation and puncture (CLP) model is the closest approximation to the mechanism of human sepsis and is now considered to be the gold standard for sepsis modeling ^[8]^. This model focuses on local necrosis caused by distal cecal ligation and systemic inflammatory response induced by leakage of intestinal content into the peritoneum. It should be noted that the severity of CLP is affected by the length of the ligated cecum, the size of the perforation needle and fluid resuscitation treatment. In addition, there are individual differences between rats, so the survival time of septic rats varies greatly. Therefore, to minimize differences in the experiment, we chose to perform ligation at half the distance between the base and the distal pole of the cecum, to keep the severity of sepsis consistent, and ZQDH decoction was administered before CLP to avoid variations in treatment.

The continuous updating of sepsis management guidelines, from the initial 6 h-bundle to 3 h-bundle, emphasizes the importance of early intervention and treatment ^[9–10]^. Therefore, compared with the observation point of 24h after CLP in most studies, we selected a relatively early observation point of 12h after CLP to simulate clinical sepsis, so as to provide a theoretical basis for early identification and rescue of clinical septic patients. According to our study, in the early stage of sepsis, the percentage of neutrophils in rats increased notably, while the levels of white blood cells and platelets decreased, prothrombin time shortened, and fibrinogen decreased, suggesting that the body was in a state of rapid depletion of clotting factors during the early inflammation outbreak, which may rapidly lead to DIC.

In traditional Chinese medicine, it is widely believed that sepsis belongs to febrile diseases, as described in ancient Chinese medicine works like *Treatise on Febrile Diseases (ShangHan Lun).* The theory of clearing heat and toxin, promoting blood circulation and removing blood stasis is now commonly used in many publications on animal and clinical sepsis ^[11]^.The prescription of ZQDH decoction consists of five herbs. Based on the theory of blood stasis and heat, Rehmanniae and Rhubarb are used as the king drugs in the prescription to clear the heat of blood division and restore the Yin of injury. Lithospermum and Radix Paeoniae Rubra are courtier drugs, which help to promote blood circulation and dissipate blood stasis, while Panax notoginseng is warm in nature as an adjuvant. These five herbs play the role of cooling blood, dispersing blood stasis, clearing heat and restoring Yin together. Studies have shown that Rhubarb has pharmacological effects such as anti-inflammation, immune regulation, endotoxin reduction, microcirculation improvement and anti-thrombosis ^[12–13]^. Among its active components, rhein and emodin have the strongest anti-inflammatory effect, while chrysophanol can inhibit platelet aggregation induced by collagen and thrombin ^[14]^. In Rehmanniae, catalpol has the effects of anti-inflammation, anti-oxidation and protection of vascular endothelium ^[15]^. Lithospermum contains rich shikonin, which can inhibit NF-κB signaling pathway, and reduce PAI-1 level as well ^[16]^. Radix Paeoniae Rubra is widely used in thrombotic diseases. Paeoniflorin is the main active ingredient of RadixPaeoniae Rubra, which not only has effects of anti-inflammation, sedation and anti-coagulation, but also protects vascular endothelial function and inhibits platelet activation by regulating the release of ET-1 and NO ^[17]^. Panax notoginseng is a classic Chinese medicine with the dual effects of hemostasis and promoting circulation. In its active components, panax notoginseng saponins have anti-inflammatory effect, dencichine can improve platelet count, and ginsenosides have effects of antithrombosis and vascular endothelial protection ^[18]^.

Platelet activation during sepsis is the main reason for platelet count decrease, the key to the formation of fibrin, and an important factor in the progression of sepsis ^[19]^. On the one hand, activated platelets participate in inflammatory reaction through aggregation, adhesion and deformation. Thus, peripheral thrombocytopenia and the increase of mean platelet volume indicate endothelial cell damage and platelet activation ^[20]^. On the other hand, activated platelets is a key regulator of blood clotting and a major source of plasma PAI-1, which plays an important role in thrombotic diseases. PAI-1, a 50 kDa glycoprotein, is the main physiological inhibitor of tissue type and urokinase type plasminogen activator (t-PA and u-PA). tPA and uPA can convert inactive plasminogen into fibrin-degrading plasminogen. Under normal circumstances, the concentration of PAI-1 in plasma and tissue remains at a low level, and only increase under pathological conditions, such as inflammation and thrombosis ^[21]^. Sepsis just consists with this pathological process. With the platelet activation in sepsis, the levels of tPA and PAI-1 in plasma and tissue increase. When PAI-1 level far exceeds tPA, fibrin cannot be degraded, leading to microthrombosis, whereas when PAI-1 level far exceeds tPA, the risk of bleeding is greatly increased.

Experimental data in this research showed that ZQDH decoction, especially at high dosage, can significantly reduce N% level, increase platelet counts and prothrombin time, and improve the infiltration of inflammatory cells in the lung, which proves that ZQDH decoction can effectively inhibit inflammation and improve coagulation function of septic rats. Simultaneously, with the platelet activation during sepsis, tPA and PAI-1 accumulate in tissues, imbalance of coagulation and fibrinolytic system leads to inhibition of fibrinolysis. Under this condition, kidney may bear the brunt of fibrin deposition and microthrombus formation. ZQDH decoction can significantly reduce septic renal fibrin deposition, and reduce renal tPA and PAI-1 levels, suggesting that ZQDH may alleviate the inhibition of fibrinolysis in sepsis by regulating the activity levels of tPA and PAI-1. However, there was no significant difference in serum tPA and PAI-1 between CLP and ZQDH groups, which may be explained by the relatively early observation window in this study, since the changes of tPA and PAI-1 in serum were consistent with those in tissues. In addition, this study found the protective effect of ZQDH decoction on lung inflammation and kidney fibrin deposition, which is also consistent with the common pathophysiological mechanisms of acute lung injury and acute kidney injury in the early stage of sepsis.

## 5. Conclusion

According to the findings obtained in this study, ZQDH decoction could inhibit systemic inflammatory injury, reduce fibrin deposition and microthrombosis, and regulate coagulation and fibrinolytic system in a dose-dependent manner. This work demonstrates the protective role of ZQDH decoction in septic rats, and opens a broad application prospect for its use in clinical practice. However, further studies are needed to explore the underlying molecular mechanisms and other key targets.

## List of abbreviations

(APTT): Activated partial thromboplastin time
(CLP): Cecal ligation and puncture
(DIC): Disseminated intravascular coagulation
(ELISA): Enzyme-linked immunosorbent assays
(FIB): Fibrinogen
(HE): Hematoxylin-eosin
(MSB): Martius-Scarlet-Blue
(MPV): Mean platelet volume
(N %): Percentage of neutrophils
(PAI-1): Plasminogen activator inhibitor-1
(PLT): Platelet count
(PT): Prothrombin time
(tPA): Tissue plasminogen activator
(uPA): Urokinase type plasminogen activator
(WBC): White blood cell count
(ZQDH): Ziqi Dihuang

